# From Bird Viremia to Bird Surveillance: Identifiability in a Multiscale Vector-Borne Model of Usutu Virus Infection

**DOI:** 10.1101/2025.11.17.688793

**Authors:** Necibe Tuncer, Yuganthi R. Liyanage, Quiyana M. Murphy, Rachel D. Persinger, Nisha K. Duggal, Stanca M. Ciupe

**Author notes:** These authors contributed equally to this work (Yuganthi R. Liyanage, Quiyana M. Murphy).

## Abstract

Usutu virus is an emerging mosquito-borne flavivirus, maintained through an enzootic cycle involving wild birds and mosquitoes, with occasional spillover to humans. Understanding how interactions across these biological scales shape transmission dynamics is essential for predicting outbreaks and improving surveillance strategies. In this study, we developed a multiscale vector-borne model of Usutu virus infection that links within-host viral kinetics in birds, the per-bite probability of mosquito infection, and population-level mosquito–bird transmission dynamics. Model parameters were validated using two laboratory datasets collected under an optimally designed experimental framework and one surveillance dataset from wild bird populations. Structural and practical identifiability analyses were conducted to evaluate parameter robustness under varying levels of measurement noise. We found that simultaneous multiscale fitting to integrated datasets improved parameter identifiability and robustness. These results highlight the importance of combining microscale and macroscale data to enhance the predictive reliability of vector-borne disease models and demonstrate the broader utility of multiscale modeling frameworks for understanding the transmission dynamics of emerging arboviruses.

**Author summary:** In this study, we developed a multiscale vector-borne model of Usutu virus infection and validated its parameters using both laboratory data collected under an optimally designed experimental framework and published surveillance data from wild bird populations. Using this model, we quantified the robustness of parameter estimates and found that multiscale fitting to integrated datasets improves the reliability and identifiability of model parameters. The results highlight the importance of combining microscale and macroscale data to enhance the predictive reliability of vector-borne disease models.

## 1 Introduction

Usutu virus is an emerging mosquito-borne arbovirus, which is maintained through an enzootic cycle involving wild birds, mosquitoes, with occasional spillover to mammals, including humans [1, 2]. Originally identified in South Africa and detected in other African countries [3], it has become endemic in most of Europe [1], affecting the avifauna and posing a risk to humans [4]. Its emergence and spread are shaped by a complex interplay of factors, including infection dynamics within-bird and within-mosquito populations, as well as virological, immunological, and environmental determinants that influence transmission to both animals and humans [4] (see (Fig. 1). Consequently, a comprehensive integrated approach that accounts for the multilevel and multiscale aspects of Usutu transmission and infection is essential to better understand Usutu virus ecology and persistence and to inform effective control strategies.

**Fig 1.**
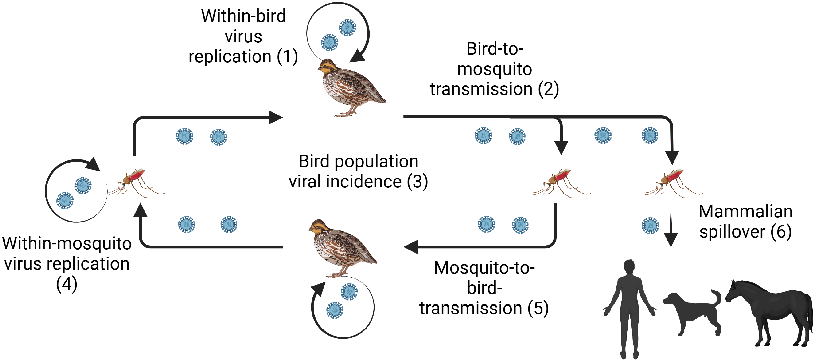
Illustration of Usutu virus enzootic transmission cycle involving several biological scales. We focus on the within-bird replication scale (1), probability of mosquito infection scale (2), and viral incidence in the bird population scale (3). Fig. created in https://BioRender.com.

We previously linked individual-level infection processes with population-level transmission dynamics by developing a multiscale mathematical model of Usutu virus infection [5]. Using this framework, we investigated how microscale variables and parameters that describe within-bird virus dynamics and the probability of mosquito infection (Fig. 1, circles 1 and 2) influence macroscale variables and parameters that govern Usutu virus incidence in bird populations (Fig. 1, circle 3). Microscale data included longitudinal measurements of viral titers in infected birds and the proportion of mosquitoes that became infected as a function of the birds’ viremia levels [6].

This multiscale model enabled a comparison between two circulating Usutu virus strains and predicted that the Netherlands 2016 Usutu strain has a higher probability of spillover to humans than the Uganda 2012 virus strain [5]. To quantify uncertainty in these predictions, we applied established structural and practical identifiability methods for ordinary differential equation models [7–9], assessed the reliability and potential biases of the resulting inferences, and proposed optimal experimental designs for non-identifiable microscale parameters. Notably, we found that virus titers should be measured every 12 hours to substantially improve model identifiability and prediction accuracy [5].

In this study, we will extend these prior results by integrating macroscale data on Usutu virus incidence in bird populations (Fig. 1, circle 3) with microscale data describing within-bird virus dynamics and the per-bite probability of mosquito infection as a function of host viremia for the Netherlands 2016 strain (Fig. 1, circles 1 and 2). The microscale data will be obtained under the previously proposed optimal experimental design, namely by collecting within-bird viral titers every 12 hours and determining the blood meal virus titers at which the majority (more than 90%) of mosquitoes become infected [5]. To determine whether the parameters of the proposed multiscale model can be uniquely estimated from ideal, noise-free data spanning all three biological scales, we will develop novel structural identifiability methods tailored for multiscale systems. In addition, to assess parameter estimability under realistic conditions of limited and noisy observations, we will conduct practical identifiability analyses.

Given the model’s integration of information across biological scales, we will investigate two complementary strategies for parameter estimation: (i) sequential estimation, in which parameters are inferred independently at each scale using scale-specific data; and (ii) simultaneous estimation, in which all parameters are estimated jointly from the entire multiscale datasets. We will compare the benefits and drawbacks of each approach and propose strategies to improve parameter estimation, model robustness, and predictive accuracy within this multiscale framework.

Ultimately, by incorporating the two-way feedback between micro- and macroscales, this work will yield a more comprehensive representation of the Usutu virus transmission cycle between birds and mosquitoes. This integrated framework will advance our understanding of Usutu virus dynamics and lifecycle and enable more accurate assessments of its zoonotic potential.

## 2 Materials and methods

### 2.1 Mathematical Model Formulation

The multiscale vector-borne model of Usutu virus infection describes the interaction between birds and mosquitoes at time *t* and the age of bird infection *τ* .

#### Model of within-bird Usutu virus dynamics

At the microscale (individual bird) level, the Usutu viral dynamics in an infected bird is modeled by a within-host acute viral infection model with an eclipse phase, as in prior work on Usutu replication in juvenile chickens [10, 11] and wild-caught house sparrows [5]. The model describes the interaction between target epithelial cells *T*, exposed epithelial cells *E*, infected epithelial cells *I*, and the Usutu virus *V*, as follows. Target cells become exposed (but not yet infectious) upon contact with the virus at rate *β* ((virus× day)^−1^). Exposed cells become infectious and start producing virus at rate *k* (day^−1^). Productively infected cells are cleared at rate *δ* (day^−1^) and produce virus at rate *π* ((cell× day)^−1^). Usutu virus is cleared at rate *c* (day^−1^). The within-host model describing these interactions is:

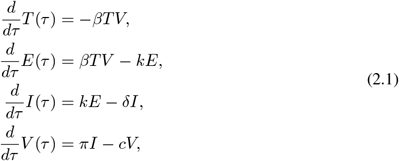

with initial condition *T* (0) = *T*_0_, *E*(0) = 0, *I*(0) = 0, *V* (0) = *V*_0_.

#### Model of per-bite probability of mosquito infection

Bird-mosquito interactions are modeled as follows. Susceptible mosquitoes become infected by biting an infected bird, with the Usutu virus transmission rate *β*_*v*_(*τ* ) = *c*_*v*_*p*_*v*_(*τ* ) (dependent on the infection age, *τ* ) being the product of the mosquitoes bite rate *c*_*v*_ ((month× bird)^−1^) and the per-bite probability of mosquito infection *p*_*v*_(*τ* ) (%). We assume that the number of Usutu viruses that are successfully transmitted from bird to mosquito at the age of infection *τ* follows a Poisson distribution with a mean transmissible virus proportional to the power of the bird’s detectable viral load (on log_10_ scale). Then, the per-bite probability of mosquito infection is:

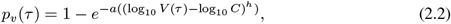

where *C* is the limit of viral detection. The form of the model Eq. (2.2) was chosen after a comprehensive model selection process [5], and *a* (virus^−1^), *C* (virus), *h* (unitless) are constant parameters [5].

#### Model of Usutu virus transmission to birds and mosquitoes

At the macroscale (bird and mosquito populations) level, we used a Ross-Macdonald-type model of vector-borne disease [12] to characterize the Usutu virus incidence in bird and mosquito populations. We assumed that mosquitoes are either susceptible to or infected with the Usutu virus. As a result, the vector population is divided into two classes: *S*_*v*_(*t*) representing the fraction of susceptible vectors at time *t* and *I*_*v*_(*t*) representing the fraction of infected vectors at time *t*. They have the same birth and death rates *µ*_*v*_ (month^−1^), to maintain a constant total population density *N*_*v*_(*t*) = *S*_*v*_(*t*) + *I*_*v*_(*t*) = 1. Similarly, birds are either susceptible to, infected with, or recovered from Usutu virus infection. To account for a bird’s infectiousness based on its viral titer, we further structured the infectious bird class with respect to infection age, *τ* . As a result, the bird population is divided into three classes: *S*_*h*_(*t*) representing the fraction of susceptible birds at time *t, i*_*h*_(*t, τ* ) representing the fraction of infected birds at time *t* with infection age *τ*, and *R*_*h*_(*t*) representing the fraction of recovered birds at time *t*. Consequently, 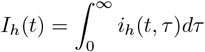 is the fraction of infected birds at time *t*, and we assumed that the total bird population is conserved *N*_*h*_(*t*) = *S*_*h*_(*t*) + *I*_*h*_(*t*) + *R*_*h*_(*t*) = 1.

We define the force of mosquito infection with respect to all infection ages *τ* as:

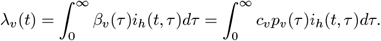

Under these assumptions, the mosquito population dynamics are described by the following differential equations:

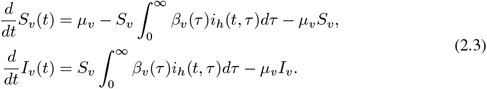

The total vector population *N*_*v*_(*t*) = *S*_*v*_(*t*) + *I*_*v*_(*t*) satisfies the differential equation 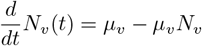, ensuring that the total vector population is asymptotically constant, 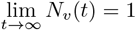.

To determine the dynamics of Usutu infection in bird populations, we defined the force of infection of susceptible birds to be:

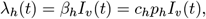

where the *β*_*h*_ ((month × mosquito)^−1^) is the Usutu virus transmission rate from infected mosquitoes to susceptible birds, *c*_*h*_ ((month × mosquito)^−1^) is the mosquito biting rate per unit time and *p*_*h*_ (%) is the per-bite mosquito-to-bird transmission probability. Both *c*_*h*_ and *p*_*h*_ are assumed to be constant. Infected birds recover at rate *γ*_*h*_ (month^−1^). Lastly, to balance the epidemic time scale (in months) and the infection age time scale (in days), we introduce a scaling parameter *κ* (day×month^−1^). Under these assumptions, the bird population dynamics are described by the following differential equations:

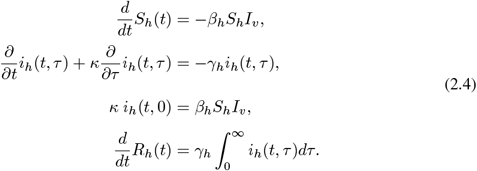

A model diagram for the multiscale model Eqs. (2.1)-(2.4) is given in Fig. 2, and a summary of its variables and parameters are given in Tables 1 and 2.

**Table 1.** Definitions and descriptions of the state variables appearing in models Eqs. (2.1)-(2.4).

| State variable | Definition |
| --- | --- |
| $S_v(t)$ | Proportion of susceptible mosquitoes at epidemic time $t$ |
| $I_v(t)$ | Proportion of infected mosquitoes at epidemic time $t$ |
| $S_h(t)$ | Proportion of susceptible birds at epidemic time $t$ |
| $i_h(t, \tau)$ | Proportion of infected birds at epidemic time $t$ and infection age $\tau$ |
| $I_h(t)$ | Proportion of infected birds at epidemic time $t$ |
| $R_h(t)$ | Proportion of recovered birds at epidemic time $t$ |
| $T(\tau)$ | Number of uninfected target cells at infection age $\tau$ |
| $E(\tau)$ | Number of exposed target cells at infection age $\tau$ |
| $I(\tau)$ | Number of productively infected target cells at infection age $\tau$ |
| $V(\tau)$ | Viral load at infection age $\tau$ |
| $p_v(\tau)$ | Per-bite probability of mosquito infection |

**Table 2.** Definitions and corresponding units of parameters in the mathematical models Eqs. (2.1)-(2.4).

|  | Parameter | Definition | Units |
| --- | --- | --- | --- |
| **Between-host** | $\beta_v(\tau)$ | Mosquito infectivity rate at bird infection age $\tau$ | $(\text{month} \times \text{bird})^{-1}$ |
| | $\mu_v$ | Mosquito birth and per capita death rate | $\text{month}^{-1}$ |
| | $\beta_h$ | Bird infectivity rate | $(\text{month} \times \text{mosquito})^{-1}$ |
| | $\gamma_h$ | Bird recovery rate | $\text{month}^{-1}$ |
| | $c_v$ | Mosquito biting rate | $(\text{month} \times \text{bird})^{-1}$ |
| | $c_h$ | Bird bitten rate | $(\text{month} \times \text{mosquito})^{-1}$ |
| **Within-host** | $\beta$ | Target cell infectivity rate | $\text{ml} \times (\text{virus} \times \text{day})^{-1}$ |
| | $k$ | Eclipse rate | $\text{day}^{-1}$ |
| | $\delta$ | Death rate of infected cells | $\text{day}^{-1}$ |
| | $\pi$ | Viral production rate | $\text{virus} \times (\text{cells} \times \text{day})^{-1}$ |
| | $c$ | Virus clearance rate | $\text{day}^{-1}$ |
| **Probability of mosquito infection** | $a$ | Positive constant | $\text{virus}^{-1}$ |
| | $h$ | Exponent | — |

**Fig 2.**
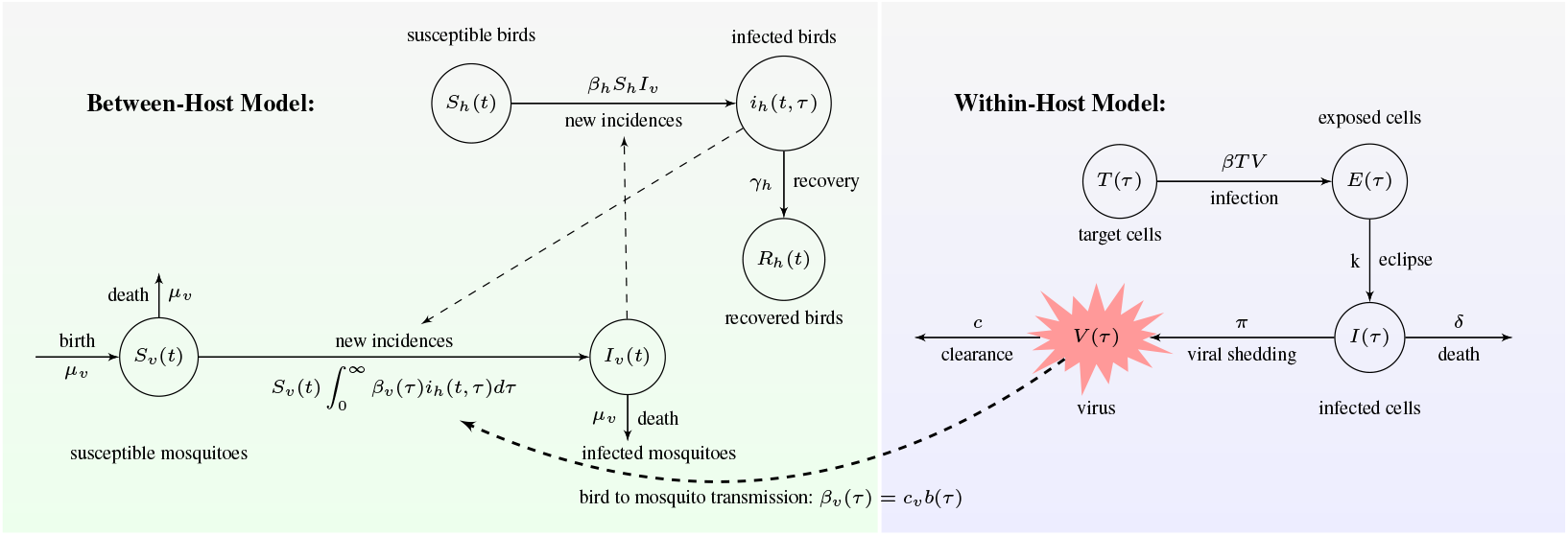
Model diagram for the between- and within-host Usutu virus infection described by Eqs. (2.1)-(2.4).

### 2.2 Parameterization and Data Fitting

#### 2.2.1 Structural Identifiability Analysis

Before validating the multiscale model Eqs. (2.1)-(2.4) with multiscale biological data, we need to determine if its parameters can be uniquely revealed given unlimited noise-free data, a process known as structural identifiability (for a review on structural identifiability, see [8, 9, 13–16]).

Model Eqs. (2.1)-(2.4) has ten variables:

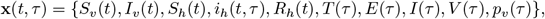

and eleven parameters:

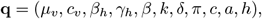

which are (initially) assumed to be unknown. To estimate parameters **q**, we will fit the model Eqs. (2.1)-(2.4) to a combination of empirical data sets consisting of viral titers in infected birds collected at the within-host time *τ* (measured in days) [17], the fraction of mosquitoes getting infected with Usutu virus when feeding on different viral titers (measured in percentage) [17], and field data on susceptible and infected birds collected at the between-host time *t* (measured in months) [18]). These measured data are noisy and consist of discrete samples of the continuous trajectory *y*(*t, τ* ). The observation *y*(*t, τ* ) is related to the model’s state variables through the mapping *g*(*x*(*t, τ* ), **q**):

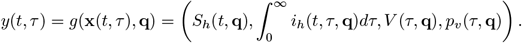

Structural identifiability aims to determine whether a unique parameter vector **q** exists for any trajectory and unlimited noise-free data *y*(*t, τ* ). If a change in any parameter in vector **q** does not affect the observed trajectory, then the parameter and (by extension) the model are structurally unidentifiable. A formal definition of structural identifiability is given below,

##### Definition 2.1. Structural Identifiability

*Let* **q**_1_ *and* **q**_2_ *be two distinct parameter vectors of the multiscale model Eqs*. (2.1)*-*(2.4). *The model is said to be structurally identifiable if and only if*

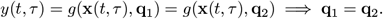

In the multiscale model given by Eqs. (2.1)–(2.4), the within-host dynamics (Eq. (2.1)) and the per-bite probability of mosquito infection (Eq. (2.2)) are embedded within the between-host transmission dynamics (Eqs. (2.3)–(2.4)). We will first perform a structural identifiability analysis of the within-host model and the per-bite probability of mosquito infection model using classical techniques [9]. Following this, we will develop a methodology to study the structural identifiability of the full multiscale system. This is a challenging task, as no standard methods currently exist for multiscale models.

#### 2.2.2 Empirical Data

##### Within-bird viral titer data

We previously demonstrated that the unknown parameters of the within-host model (Eq. (2.1)) are practically unidentifiable when viral titers are measured only once per day following bird inoculation. Through *in-silico* experiments, we predicted that this parameter unidentifiability could be resolved if viral titers were collected every 12 hours for up to seven days post-inoculation [5]. Based on this theoretical result, we designed an inoculation study consisting of two cohorts of canary birds [17]. The first cohort included 13 birds, and the second cohort included 10 birds. Each bird was inoculated with 1500 plaque-forming units (PFU) of Usutu Netherlands 2016 virus strain. Infectious viral titers (measured in PFU ml^−1^) were collected daily post-inoculation from day 1 to day 6 for the first cohort *τ* ^1^ = {1, 2, 3, 4, 5, 6} days, and from day 0.5 to day 6.5 for cohort 2 *τ* ^2^ = {0.5, 1.5, 2.5, 3.5, 4.5, 5.5, 6.5} days [17]. Three birds died during the study, and all remaining birds exhibited undetectable viremia by day 6 post-inoculation; therefore, data collected at day 6.5 were excluded from further analyses. We averaged virus titers among all birds at time points *τ*_*w*_ = {0.5, 1, …, 5.5, 6} days, thus creating a population-level data set *V* ^*data*^(*τ*_*w*_) of Usutu virus titers collected twice a day (see Table 3).

**Table 3.** Longitudinal Usutu virus (Netherlands 2016 strain) titers (log_10_ PFU/ml) measured in infected canaries over time (days post–inoculation).

| Time (days) | 0.5 | 1 | 1.5 | 2 | 2.5 | 3 | 3.5 | 4 | 4.5 | 5 | 5.5 | 6 |
| --- | --- | --- | --- | --- | --- | --- | --- | --- | --- | --- | --- | --- |
| $\log_{10}$ Virus<br>($\log_{10}$ PFU/ml) | 1.7 | 2.65 | 2.68 | 5.2 | 4.25 | 6.46 | 5.0 | 4.15 | 4.2 | 2.72 | 3.0 | 1.7 |

##### Data on percent mosquito infection based on viral load exposure

We collected data on the percentage of mosquito infection as a function of viral load exposure, as follows. Nine cartons of *Culex pipiens* mosquitoes (ranging from 26 to 47 individuals per carton) were fed on cotton balls soaked with nine distinct concentrations of Usutu Netherlands 2016 virus strain spanning increasing magnitudes. At 10 days post–blood meal, mosquitoes were tested for Usutu virus infection and classified as infected if viral RNA was detected in their body. The resulting data on percent mosquito infection as a function of virus load exposure are summarized in Table 4 [17].

**Table 4.** Percent of *Culex pipiens* mosquitoes infected after feeding on blood meals spiked with varying Usutu virus (Netherlands 2016 strain) concentrations, expressed in plaque-forming units per milliliter (PFU/ml).

|  |  |  |  |  |  |  |  |  |  |
| --- | --- | --- | --- | --- | --- | --- | --- | --- | --- |
| $\log_{10}$ Virus<br>(PFU/ml) | 3.5 | 4.2 | 5.3 | 5.4 | 6.3 | 6.4 | 6.5 | 7.4 | 7.9 |
| Percent infection<br>($\times 100$) | 0 | 0 | 5.9 | 0 | 16.7 | 36.7 | 29.4 | 73.1 | 93.6 |

##### Data on Usutu virus incidence in wild birds

We used previously published passive surveillance data on Usutu virus incidence in dead birds collected at a wildlife rehabilitation center in Ferrara (Italy), where mosquitoes and wild birds co-exist, and Usutu virus has become endemic [18, 19]. Usutu virus incidence was recorded twice per month from 2015 to 2019 [18]. The bimonthly incidence data, aggregated across the five-year period, indicated that bird infections occurred between June 16 and November 15, with peak infection observed in the latter half of August (see Table 5).

**Table 5.** Proportions of birds tested (susceptible) and infected with Usutu virus between June 16 and November 15, aggregated over five years. Data were digitized from Fig. 1 in [18] using the *Grabit* tool in MATLAB. In [18], the number of birds tested and infected are reported. The susceptible population was converted to proportions by dividing each sample by the maximum observed sample size.

| Time<br>(months) | 0 | 0.5 | 1 | 1.5 | 2 | 2.5 | 3 | 3.5 | 4 | 4.5 |
| --- | --- | --- | --- | --- | --- | --- | --- | --- | --- | --- |
| Fraction of Susceptible<br>Bird Population | 0.99 | 0.61 | 0.73 | 0.62 | 0.43 | 0.29 | 0.29 | 0.24 | 0.25 | 0.18 |
| Fraction of Infected<br>Bird Population | 0.0066 | 0.038 | 0.136 | 0.152 | 0.219 | 0.095 | 0.006 | 0.0197 | 0.032 | 0 |

Data availability at three biological scales allows flexibility in parameter estimation. Because the within-host model (Eq. (2.1)) and the per-bite mosquito infection model (Eq. (2.2)) are independent of the between-host model parameters (Eqs. (2.3)–(2.4)), we can adopt a sequential fitting approach, estimating microscale parameters first and fixing them before fitting the macroscale model [15, 20]. Alternatively, a simultaneous data fitting strategy can be employed, where all parameters of the full multiscale model (Eqs. (2.1)–(2.4)) are estimated jointly using the complete multiscale datasets [15, 20].

#### 2.2.3 Sequential Data Fitting

##### Fitting the within-host model

We assumed that all initial conditions for the within-host model (Eq. (2.1)) are known to ensure its structural identifiability (see Proposition 3.3). Specifically, we set the target epithelial cells to *T* (0) = 4 × 10^6^ cells/ml; assume that there are no exposed or infected cells at the start of infection, *E*(0) = *I*(0) = 0 cells/ml; and set the initial viral titer to *V* (0) = 1 PFU/ml, as in [5]. The parameters **p**_**1**_ = {*β, k, δ, c, π*} of the within-host model (Eq. (2.1)), all of which are treated as unknown, are estimated by minimizing the least-squares functional:

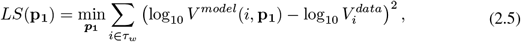

where *V* ^*model*^(*i*, **p**_**1**_) is Eq. (2.1)-predicted viral load at time *i* (on a log_10_ scale) and 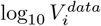 is the observed average virus titers (also on a log_10_ scale) at time points *i* ∈ *τ*_*w*_ = {0.5, 1, 1.5, …, 6} days post infection (see Table 3). Parameter estimation was performed using MATLAB’s *fmincon* optimization algorithm, subject to parameter bounds *β* ∈ [10^−10^, 7.4], *k* ∈ [0.5, 7], *δ* ∈ [0, 25], *π* ∈ [0, 10^8^], and *c* ∈ [0, 50].

##### Fitting the probability of mosquito infection model

The per-bite probability of mosquito infection curve (Eq. (2.2)) is structurally identifiable (see Proposition 3.4). Its parameters ***p***_**2**_ = {*a, h*}, which are assumed unknown, are estimated by minimizing the least-squares functional:

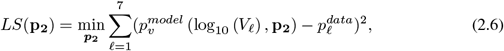

where 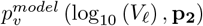 is Eq. (2.2)-predicted per-bite probability of mosquito infection with log_10_ *C* = 3.5. log_10_ *V*_*ℓ*_ are viral titers in the artificial meal at the day of mosquito feed, and 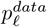 are the percentage of infected mosquitoes, ten days after the artificial blood meal. (see Table 4). Parameter estimation was performed using MATLAB’s *fmincon* optimization algorithm, subject to parameter bounds *a* ∈ [10^−5^, 10^−2^] and *h* ∈ [0, 10].

##### Fitting the between-host model

We assume that at the beginning of the outbreak *I*_*v*_(0) = 0.01 mosquitoes are infected and *S*_*v*_(0) = 0.99 mosquitoes are susceptible. Additionally, *i*_*h*_(0, *τ* ) = *i*_0_(*τ* ) = 4.2 × 10^−7^ birds are infected in each age of infection *τ* ; no birds have recovered *R*_*h*_(0) = 0; and the remaining susceptible bird population is 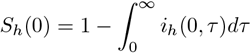. The parameters ***p***_**3**_ = {*β*_*h*_, *c*_*v*_, *γ*_*h*_, *µ*_*v*_} of the between-host model (Eqs. (2.3)-(2.4)), which are assumed unknown, are estimated by minimizing the least-squares functional:

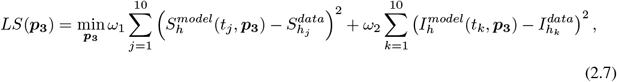

where 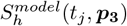 and 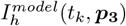 represent the model-simulated proportions of susceptible and infected birds at time points *t*_*j*_ and *t*_*k*_; 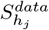 and 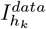 are empirical data on proportions of susceptible and infected birds (see Table 5); and *ω*_1_ = 1, *ω*_2_ = 10 are weights that scale the two populations.

Parameter estimation was performed using the MATLAB’s *fmincon* optimization algorithm combined with the *MultiStart* function with 1000 initial guesses, subject to parameter bounds *β*_*h*_ ∈ [0, 10^2^], *c*_*v*_ ∈ [0, 10^3^], *γ*_*h*_ ∈ [1*/*7, 1] (to account for bird recovery periods between 1 and 7 months), and *µ*_*v*_ ∈ [0.5, 6] (to account for female mosquito average lifespan between 5 days and 2 months [21]).

#### 2.2.4 Simultaneous Data Fitting

For the simultaneous data fitting procedure, we assumed that the parameters of the per-bite probability of mosquito infection model (Eq. (2.2)) are known, *a* = 1.6 × 10^−3^ virus^−1^ and *h* = 4.97 (see Table 7). We estimate the remaining parameters **p** = {*β*_*h*_, *c*_*v*_, *γ*_*h*_, *µ*_*v*_, *β, k, δ, π, c*} (**p**_**1**_ = {*β, k, δ, π, c*} for the within-host model and **p**_**3**_ = {*β*_*h*_, *c*_*v*_, *γ*_*h*_, *µ*_*v*_} for the between-host model) by fitting the multiscale model (Eq. (2.1)-(2.4)) to the combined viral titers in infected birds data (Table 3) and the proportions of susceptible and infected bird data (Table 5).

**Table 6.** Parameter estimates obtained by fitting the within-host model Eq. (2.1) to virus titer data in Table 3.

| Parameter | Definition | Value | Bounds | Units |
| --- | --- | --- | --- | --- |
| $\beta$ | Target cell infectivity rate | $10^{-5}$ | $[10^{-10}, 7.4]$ | $\text{ml} \times (\text{PFU} \times \text{day})^{-1}$ |
| $k$ | Eclipse phase | 6.99 | $[0.5, 7]$ | $\text{day}^{-1}$ |
| $\delta$ | Infected cells death rate | 19.5 | $[0, 25]$ | $\text{day}^{-1}$ |
| $\pi$ | Virus production rate | 10.52 | $[0, 10^8]$ | $\text{PFU} \times (\text{cell} \times \text{day})^{-1}$ |
| $c$ | Virus clearance rate | 3.49 | $[0, 50]$ | $\text{day}^{-1}$ |

**Table 7.** Parameter estimates obtained by fitting the per-bite probability of mosquito infection *p*_*v*_(*τ* ) given by Eq. (2.2) to the percent of mosquito infection data in Table 4.

| Parameter | $a$ | $h$ |
| --- | --- | --- |
| Bounds | $[10^{-5}, 10^{-2}]$ | $[0, 10]$ |
| Value | $1.6 \times 10^{-3}$ | 4.97 |
| AIC | -61.59 |  |

We estimate parameters **p** by minimizing the following least squares functional:

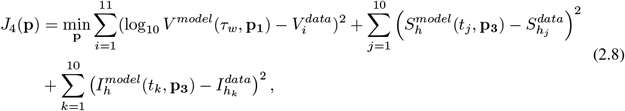

using the MATLAB’s *fmincon* optimization algorithm combined with the *MultiStart* function with 1000 initial guesses, subject to parameter bounds *β*_*h*_ ∈ [0, 100], *c*_*v*_ ∈ [0, 10^3^], *γ*_*h*_ ∈ [1*/*7, 1], *µ*_*v*_ ∈ [0.5, 6],*β* ∈ [10^−10^, 7.4], *k* ∈ [0.5, 7], *δ* ∈ [0, 25], *π* ∈ [0, 10^8^], and *c* ∈ [0, 50] (as in the sequential data fitting algorithm).

#### 2.2.5 Practical Identifiability Analysis

Structural identifiability assesses whether parameters can be uniquely determined from a model with the ideal condition of unlimited, noise-free data. However, in practice, data are often discrete and contaminated with measurement errors, making structural identifiability insufficient to guarantee practical identifiability. Practical identifiability determines whether parameters can be accurately estimated under such conditions. Several techniques are available for assessing the practical identifiability of ODE models [8, 9, 16, 22, 23]. In this study, we use the Monte Carlo simulation (MCS) approach [8, 9, 16], which involves the following steps:

1. We numerically solve the considered model for best parameter estimates **p**, obtained by fitting it to the discrete experimental data. For the within-host model, we solve Eq. (2.1) numerically with the estimated parameter **p**_**1**_ to obtain the model predictions log_10_ *V* ^*model*^(*τ*_*w*_, **p**_**1**_) at the experimental time points *τ*_*w*_. For the per-bite probability of mosquito infection model, we solve Eq. (2.2) numerically with the estimated parameter **p**_**2**_ to obtain the model predictions 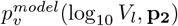. For the between-host model, we solve Eqs. (2.3)-(2.4) numerically with the estimated parameter **p**_**3**_ to obtain the model predictions 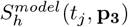 and 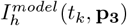.
2. We generate *M* = 1000 synthetic datasets using the following constant error models by adding *σ* = {1%, 5%, 10%, 20%} measurement errors to each given experimental data point. The error models are as follows: <u>Within-host scale</u>:

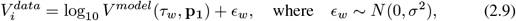

<u>Probability of mosquito infection scale:</u>

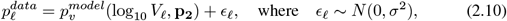

<u>Between-host scale:</u>

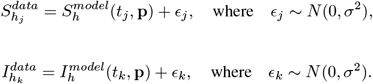

This constant error model does not always preserve the non-negativity of the synthetically generated, noise-perturbed infected bird population data 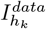. To ensure positivity, we sample the synthetic data from a log-normal distribution, using the following multiplicatie error model:

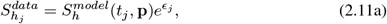

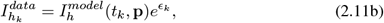

while keeping the viral titer data as before.
3. For each measurement error *σ*, we fit each model, sequentially or simultaneously, to each of the 1000 generated datasets to estimate new parameter values **q**_*i*_, where *i* = 1, 2, …, 1000.
4. We calculate the average relative estimation error (ARE) for each parameter of the chosen model at each measurement error *σ*

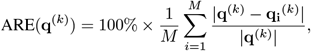

where **q**^(*k*)^ is the k-th element of the parameter set **q** and **q**_**i**_^(*k*)^ is k-th element of **q**_**i**_

We use the ARE formula to assess the practical identifiability of each parameter in the model, according to Definition 2.2 provided below [24]:

##### Definition 2.2.

*The practical identifiability of a parameter* **q** *is determined by comparing its average relative estimation error ARE with the measurement error σ. We say that the parameter* **q** *is:*

i. *Strongly practically identifiable if ARE*(**q**) ≤ *σ*,
ii. *Weakly practically identifiable if σ < ARE*(**q**) ≤ 10*σ*,
iii. *Practically unidentifiable if ARE*(**q**) *>* 10*σ*.

*A model is said to be practically identifiable when all its parameters are practically identifiable*.

## 3 Results

### 3.1 Structural Identifiability Results

#### Structural identifiability at the within-host scale

For model Eq. (2.1), the available data consist of longitudinal viral titers collected after bird inoculation with the Netherlands 2016 Usutu virus strain. The structural identifiability of ODE models, including Eq. (2.1), is well established in the literature, and various methods exist to determine whether parameters can be uniquely identified from observations [24]. In our previous study, we assessed the structural identifiability of Eq. (2.1) using the differential algebra method [5]. This approach rewrites Eq. (2.1) using only the observed variable *V* (*τ* ) and model parameters, yielding the input-output differential polynomial:

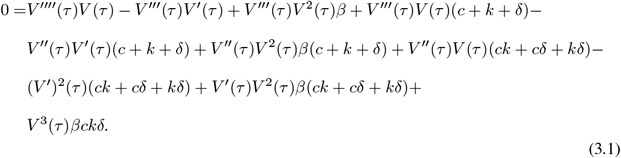

Structural identifiability is then determined by checking whether the mapping from parameters to the coefficients of the input-output equation Eq. (3.1) is one-to-one. Formally, we have:

##### Definition 3.1.

*Let c*(**q**) *denote the coefficients of the input-output equation Eq*. (3.1). *We say that the within-host model (Eq*. (2.1)*) is globally structurally identifiable given unlimited observations for the virus population V* (*τ* ) *if and only if*,

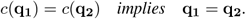

Applying Definition 3.1, we set *c*(**q**_**1**_) = *c*(**q**_**2**_) and obtain the following system of nonlinear equations:

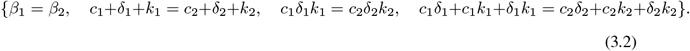

From Eq. (3.2), only the infection rate *β* can be uniquely determined. The viral clearance rate *c*, infected cell death rate *δ*, and eclipse rate *k* are interchangeable, resulting in six possible solutions [5], so they are locally identifiable. The viral production rate *π* does not appear in Eq. (3.2), meaning infinitely many values of *π* yield the same observations; hence *π* is unidentifiable.

##### Proposition 3.2.

*The within-host model (Eq*. (2.1)*) with unknown initial conditions is not structured to identify all its parameters from unlimited viral load observations V* (*τ* ).

*Specifically, β is globally structurally identifiable, c, δ, k are locally structurally identifiable, and π is unidentifiable*.

A structurally unidentifiable model can be transformed into a structurally identifiable one by fixing unidentifiable parameters or initial conditions. Here, by fixing the initial conditions of Eq. (2.1), the model becomes fully identifiable [5]:

##### Proposition 3.3.

*The within-host model (Eq*. (2.1)*) with known initial conditions is structured to identify all its parameters from unlimited, noise-free viral load observations V* (*τ* ).

#### Structural identifiability at the per-bite probability of mosquito infection scale

For model Eq. (2.2), data consist of the fraction of mosquitoes infected after feeding on blood containing different viral loads. As before, structural identifiability involves determining whether the mapping from the parameter space to the observations is one-to-one. By setting the model outputs for two sets of distinct parameters equal to each other, we obtain:

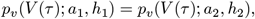

which is equivalent to:

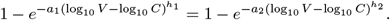

This equality implies that

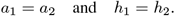

##### Proposition 3.4.

*The per-bite mosquito infection probability (Eq*. (2.2)*) is structurally identifiable given unlimited, noise-free observations of the fraction of mosquitoes infected after feeding on blood containing continuous virus loads*.

#### Structural identifiability at the between-host scale

The between-host model Eqs. (2.3)-(2.4) encompasses nested models Eq. (2.1) and Eq. (2.2), making its identifiability analysis inherently multiscale. As with the microscale models, we determine structural identifiability of Eqs. (2.3)-(2.4) by checking whether the mapping from the parameter space into the observations is one-to-one. This requires first deriving input-output equations consisting of only the observed variables. By solving model Eq. (2.4) using the method of characteristics we obtain:

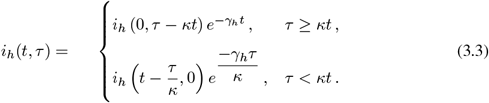

Substituting the boundary condition *κi*_*h*_(*t*, 0) = *β*_*h*_*S*_*h*_*I*_*h*_ and initial condition *i*_*h*_(0, *τ* ) = *i*_0_(*τ* ), we obtain:

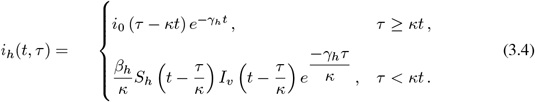

As mentioned before, we will calibrate the model using only field data for susceptible and infected bird populations, *S*_*h*_ (*t*) and 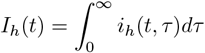, since data for the (susceptible and infectious) vector populations is not available. Thus, to obtain input-output equations for observed variables, we must eliminate the mosquito populations from Eqs. (2.3)-(2.4).

Assuming the mosquito population is asymptotically constant, *S*_*v*_(*t*) + *I*_*v*_(*t*) = 1, the infected vector population can be rewritten as [25]:

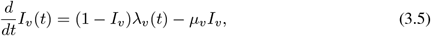

where *λ*_*v*_(*t*) is the force-of-infection within vector population. Next, by integrating the infected bird state Eq. (2.4) with respect to infection age *τ*, we obtain:

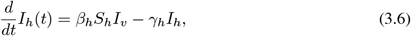

where we used the fact that 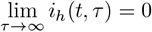. Finally, solving for *I*_*v*_ from Eq. (3.6) and substituting into Eq. (3.5) yields an input-output equation relating only the observed bird populations *S*_*h*_(*t*) and *I*_*h*_(*t*) to the model parameters:

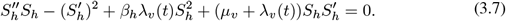

We can expand Eq. (3.7), by substituting the force of infection *λ*_*v*_(*t*) and the infected bird population *i*_*h*_(*t, τ* ) from Eq. (3.4), to obtain an explicit input-output equation:

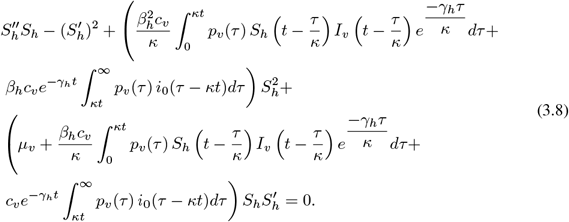

By substituting 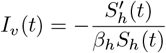 into Eq. (3.6), we obtain a differential relationship between the observed bird populations:

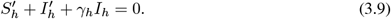

At the between-host scale, we thus derived two input-output equations: Eqs. (3.8) and (3.9). Eq. (3.9) is a differential polynomial, analogous to those obtained in the ODE-based within-host models. In contrast, Eq. (3.8) is a differential-integro polynomial, involving both derivatives and integrals of the observable variables. Such integral terms naturally arise because they represent the cumulative nature of infection in between-host dynamics. The complete list of input-output equations for the multiscale model Eqs. (2.1)-(2.4) becomes:

**<u>Between-host scale</u>:**

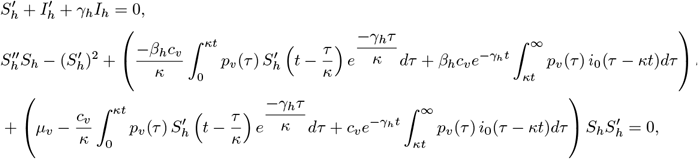

**<u>Per-bite probability of mosquito infection scale:</u>**

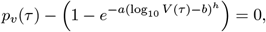

**<u>Within-bird scale:</u>**

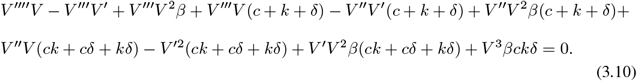

##### Theorem 3.5.

*The multiscale model Eqs*. (2.1)*–*(2.4) *is structurally identifiable if unlimited, noise-free observations are available for: (i) viral load within an infected bird, (ii) the fraction of mosquitoes infected when feeding on blood with varying virus loads, and (iii) field data on susceptible and infected birds, provided that the initial infected-bird distribution i*_0_(*τ* ) *and the initial conditions of Eq*. (2.1) *are known. In the absence of these conditions, the multiscale model is unidentifiable*.

*Proof*. For the multiscale model to be structurally identifiable, we must show that the mapping from the parameter space to the available observations is one-to-one. To simplify the notation, we define:

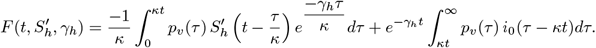

Then the input-output equations (Eqs. (3.10)) for the multiscale model simplify to:

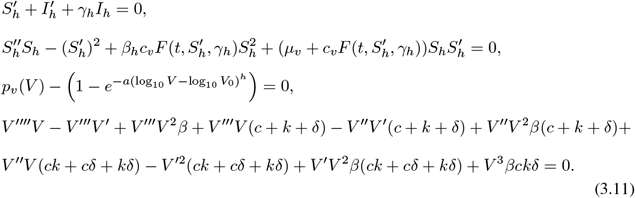

As before, we consider two sets of parameters:

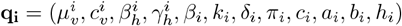, for *i* = 1, 2 and set the coefficients of the input-output equations (Eqs. (3.11)) equal to each other, i.e. *c*(**q**_**1**_) = *c*(**q**_**2**_). This results in the following system of nonlinear equations:

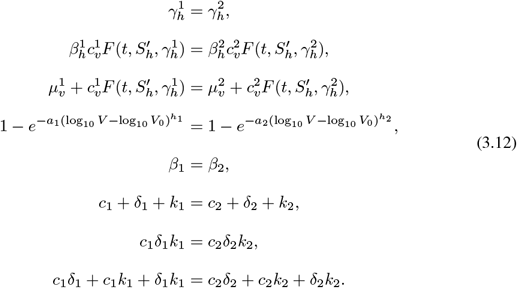

By propositions (3.3) and (3.4), the last five equations in Eq. (3.12) admit unique solutions, implying that the parameters *a, h, β, k, δ, π*, and *c* are structurally identifiable. Moreover, since 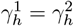, it follows that:

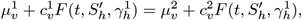

which gives 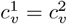 and 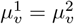. Furthermore, from:

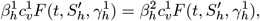

we obtain 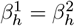. Hence, the multiscale model Eqs. ((2.1)-(2.4)) is identifiable when the within-host model (Eq. (2.1)) is identifiable (which is guaranteed by known initial conditions) and unidentifiable otherwise. □

### 3.2 Data Fitting Results

#### 3.2.1 Sequential Data Fitting

##### Within-host results

Parameters of the within-host model (Eq. (2.1)) were estimated by minimizing the distance between the predicted viral curve (on log_10_ scale) and the average virus titers data (also on log_10_ scale) from birds inoculated with the Netherlands 2016 Usutu virus strain given in Table 3. The best fit parameter estimates for within-host parameters **p**_**1**_ = {*β, k, δ, π, c*} are presented in Table 6. The predicted virus curve and the 95% confidence region are presented in Fig. 3.

**Fig 3.**
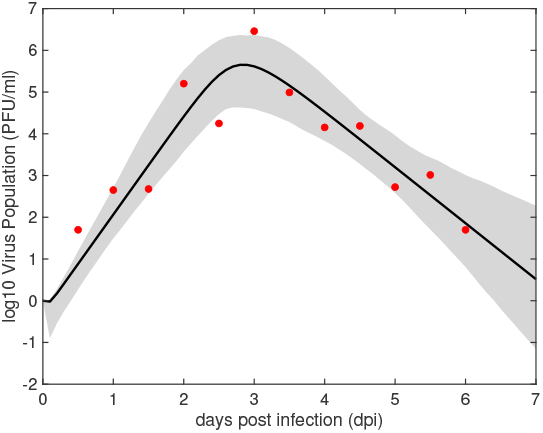
Virus dynamics (solid line) and 95% confidence region (shaded area) obtained by fitting *log*_10_ *V* given by the within-host model (Eq. (2.1)) to virus titer data (red dots). The estimated parameters are given Table 6.

Model Eq. (2.1) predicts fast virus expansion to peak concentration of 6.7 × 10^5^ PFU/ml at 2.8 days post inoculation. This is followed by decays below 10 PFU/ml after day 6 post inoculation (Fig. 3, black curve). We observe large variability around the predicted curve, with the widest confidence region occurring around the decay region (Fig. 3, shaded area). The estimated infectivity rate is *β* = 10^−5^ ml × (PFU × day)^−1^, the eclipse phase is *k* = 6.99 day^−1^ (corresponding to an exposed cell lifespan of 3.4 hours), and the infected cell death rate is *δ* = 19.5 day^−1^ (corresponding to an infected cell lifespan of 1.2 hours). Additionally, the estimated virus production rate is *π* = 10.52 PFU × (cell × day)^−1^ and the virus clearance rate is *c* = 3.49 day^−1^ (corresponding to a virion lifespan of 6.8 hours).

We compared these results to parameter estimates from a within-host Usutu virus infection in wild-caught house sparrows inoculated with the same Netherlands 2016 strain [5]. Besides the host species, one notable difference between the studies was the sampling frequency: viral titers in house sparrows were measured once daily for up to seven days, whereas in canaries, samples were collected twice daily for up to 6.5 days. We investigated whether this difference in sampling frequency affected parameter estimates.

Our analysis revealed identical estimates for target cell infectivity rates *β* and similar estimates for the eclipse rate *k*. The remaining parameters differ among the two studies, with the virus production rate *π* and infected cell death rate *δ* being 1.4-times and 2.8-times larger in canaries than in house sparrows, *π* = 10.52 PFU× (cell × day)^−1^ and *δ* = 19.5 day^−1^ versus *π* = 7.49 (cell × day)^−1^ and *δ* = 6.95 day^−1^, respectively. The largest difference is in the viral clearance rate *c* estimate, which is 14-times smaller in canaries than in house sparrows, *c* = 3.49 day^−1^ versus *c* = 48.8 day^−1^. Consequently, the within-host basic reproduction number was higher in canaries (ℛ_0_ = 6.18) compared to house sparrows (ℛ_0_ = 4.2).

##### Per-bite probability of mosquito infection results

We estimated parameters **p**_**2**_ = {*a, h*} by minimizing the distance between the predicted per-bite probability of mosquito infection (Eq. (2.2)) and the percent of mosquitoes getting infected when feeding on blood meals containing Usutu virus (see Table 4). The best fit parameter estimates are given in Table 7. The predicted probability of mosquito infection curve and 95% prediction region are presented in Fig. 4. We predicted that *a* = 1.6 × 10^−3^ (log_10_PFU)^−1^ and *h* = 4.97 (unitless). This resulted in a per-bite probability of mosquito infection that follows a sigmoidal power-law shape, with an inflection point at 10^7^ PFU/ml (see Fig. 4, black curve). There is limited variability in the results with tight 95% confidence levels (see Fig. 4, shaded area).

**Fig 4.**
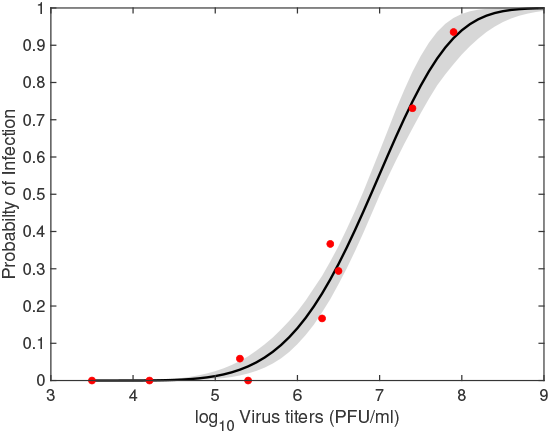
Predicted per-bite probability of mosquito infection (solid line) and 95% confidence region (shaded region) obtained by fitting model Eq. (2.2) to data (red dots). The estimated parameters are given Table 7.

##### Between-host scale results

We fixed the within-host and the per-bite probability of mosquito infection parameters (Eqs. (2.1)-(2.2)) at their estimated values given in Tables 6 and 7, then estimated the between-host parameters ***p***_**3**_ = {*β*_*h*_, *c*_*v*_, *γ*_*h*_, *µ*_*v*_} by fitting the model Eqs. (2.3)-(2.4) to proportions of susceptible and infected birds given in Table 5. The resulting best parameter estimates are provided in Table 8. The predicted susceptible and infected bird curves and their 95% prediction region are presented in Fig. 4.

**Table 8.** Parameter estimates obtained by fitting the between-host model (Eqs. (2.3)-(2.4)) to proportions of susceptible and infected birds given in Table 5.

| Parameter | Description | Value | Bounds | Units |
| --- | --- | --- | --- | --- |
| $\beta_h$ | Bird infectivity rate | 6.86 | $[0, 100]$ | (month × mosquito)$^{-1}$ |
| $c_v$ | Mosquito biting rate | 338.71 | $[0, 10^3]$ | (month × bird)$^{-1}$ |
| $\gamma_h$ | Bird recovery rate | 0.14 | $[1/7, 1]$ | month$^{-1}$ |
| $\mu_v$ | Mosquito birth/death rate | 2.87 | $[0.5, 6]$ | month$^{-1}$ |

Model Eqs. (2.3)–(2.4) predict an epidemic peak involving approximately 11% of the bird population occurring 1.58 months post-exposure (Fig. 5, right panel, black curve). The epidemic subsequently declines, with fewer than 0.1% of birds remaining infected by 5 months post-exposure, at which point 84% of the population has recovered (Fig. 5, left panel, black curve). Model uncertainty is low, as indicated by the narrow 95% confidence intervals (Fig. 5, shaded region). We predict that the bird population infectivity rate is *β*_*h*_ = 6.86 (month × mosquito)^−1^, the mosquito biting rate is *c*_*v*_ = 338.71 (month × bird)^−1^, the mosquito birth/ death rate is *µ*_*v*_ = 2.87 month^−1^ (corresponding to a mosquito lifespan of 0.35 months) [26], and the recovery rate is *γ*_*h*_ = 0.14 month^−1^ (corresponding to an epidemic of seven months, similar to the reported data [18]).

**Fig 5.**
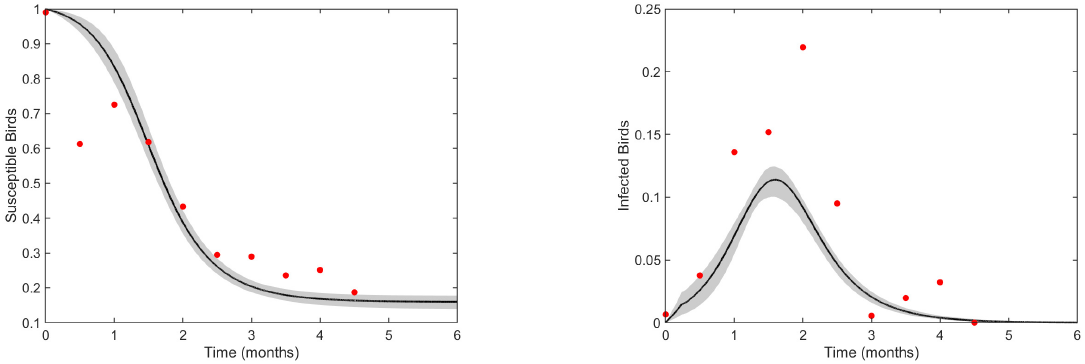
Predicted susceptible (left) and infected (right) bird percentages (solid lines) and 95% confidence intervals (shaded area) obtained from fitting the between-host model (Eqs. (2.3)-(2.4)) to field data (red dots). Estimated parameter values are given in Table 8.

#### 3.2.2 Simultaneous Data Fitting

The eleven parameters **p** = {*a, h, β*_*h*_, *c*_*v*_, *γ*_*h*_, *µ*_*v*_, *β, k, δ, π, c*} of the multiscale model (Eqs. (2.1)-(2.4)) can all be identified from unlimited noise-free observations of within-host viral load, the percent of mosquito infection, and the proportions of susceptible and infected birds when the initial values of the within-host model (Eq. (2.1)) are known (see Theorem 3.5 for details). We fixed parameters *a* and *h* in Eq. (2.2) to the values in Table 7 and estimated the remaining nine parameters by simultaneously fitting Eq. (2.1) and Eqs. (2.3)-(2.4) to viral load data in Table 3 and surveillance susceptible and infected birds data in Table 5. The best fit parameters obtained by minimizing the least squares functional Eq. 2.8 are presented in Table 9. The predicted viral load, proportion of susceptible, and proportion of infected birds and their 95% prediction regions are presented in Fig. 6.

**Table 9.** Parameter estimates obtained by fitting the the multiscale model (Eqs. (2.1)-(2.4)) to viral titer data in Table 3 and the proportions of susceptible and infected birds given in Table 5.

| Parameter | Description | Value | Bounds | Units |
| --- | --- | --- | --- | --- |
| $\beta_h$ | Bird infectivity rate | 8.47 | $[0, 100]$ | $(\text{month} \times \text{mosquito})^{-1}$ |
| $c_v$ | Mosquito biting rate | 548.28 | $[0, 10^3]$ | $(\text{month} \times \text{bird})^{-1}$ |
| $\gamma_h$ | Bird recovery rate | 0.14 | $[1/7, 1]$ | $\text{month}^{-1}$ |
| $\mu_v$ | Mosquito birth/death rate | 4.97 | $[0.5, 6]$ | $\text{month}^{-1}$ |
| $\beta$ | Target cell infectivity rate | $1.28 \times 10^{-5}$ | $[10^{-10}, 7.38]$ | $\text{ml} \times (\text{PFU} \times \text{day})^{-1}$ |
| $k$ | Eclipse phase | 6.99 | $[0.5, 7]$ | $\text{day}^{-1}$ |
| $\delta$ | Infected cells death rate | 23.72 | $[0, 25]$ | $\text{day}^{-1}$ |
| $\pi$ | Virus production rate | 8.73 | $[0, 10^8]$ | $\text{PFU} \times (\text{cells} \times \text{day})^{-1}$ |
| $c$ | Virus clearance rate | 3.22 | $[0, 50]$ | $\text{day}^{-1}$ |

**Fig 6.**
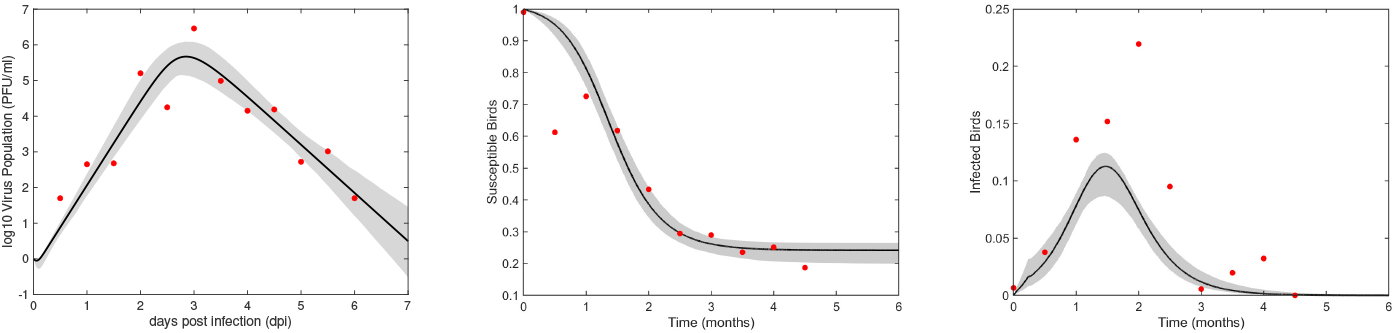
Predicted virus (left), susceptible bids (middle) and infected birds (right) (solid lines), and 95% confidence intervals (shaded area) obtained from fitting the multiscale model (Eqs. (2.1)-(2.4)) to data (red dots). Estimated parameter values are given in Table 9.

We observe no differences in the peak virus dynamics or the size of the viral load at the end of the experiment between sequential and simultaneous data fitting results (see Fig. 3 versus Fig. 6), although sequential data fitting yields tighter predictions. By contrast, we predict differences in both the magnitude and timing of the epidemic peak, as well as in the final epidemic size (by 9% and 33%, respectively), between sequential and simultaneous data fitting results (see Fig. 5 versus Fig. 6). These discrepancies are primarily driven by larger differences in parameter estimates. Specifically, when comparing within-host parameter estimates obtained through sequential versus simultaneous data fitting, we observe notable differences in the average host cell infectivity rate (*β*), the average infected cell death rate (*δ*), and the average virus production rate (*π*). Estimates from the simultaneous data fitting yielded a higher average host cell infectivity rate, *β* = 1.28 × 10^−5^ ml × (PFU × day)^−1^, compared to *β* = 1.0 × 10^−5^ ml × (PFU × day)^−1^ obtained from sequential data fitting. The average lifespan of infected cells was slightly shorter under simultaneous fitting, with 1*/δ* = 0.042 days (approximately 1 hour), compared to 1*/δ* = 0.051 days (about 1.2 hours) for the sequential fitting scenario. In contrast, the average virus production rate was lower under simultaneous fitting, *π* = 8.73 PFU × (cells × day)^−1^, representing approximately 0.83 times the value estimated from sequential fitting (*π* = 10.52 PFU × (cells × day)^−1^). The differences in the average eclipse rate (*k*) and virus clearance rate (*c*) were negligible between the two fitting approaches (see Table 9 and Table 6). When comparing between-host parameter estimates obtained through sequential and simultaneous data fitting, we observe differences in the mosquito biting rate (*c*_*v*_), bird infectivity rate (*β*_*h*_), and mosquito birth/death rate (*µ*_*v*_). Simultaneous data fitting resulted in higher estimates for both the mosquito biting rate, *c*_*v*_ = 548 (month × bird)^−1^ compared to *c*_*v*_ = 338.71 (month × bird)^−1^ from sequential data fitting, and the bird infectivity rate, *β*_*h*_ = 8.47 (month × mosquito)^−1^ compared to *β*_*h*_ = 6.86 (month × mosquito)^−1^ in the sequential fitting. In contrast, the estimated mosquito lifespan was shorter under simultaneous fitting, with 1*/µ*_*v*_ = 0.2 months (approximately 6 days) compared to 1*/µ*_*v*_ = 0.34 months (about 10.2 days) in the sequential fitting scenario (see Table 9 and Table 8).

Lastly, we define the probability that a bird bitten by an infected mosquito acquires Usutu virus infection to be:

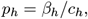

where *β*_*h*_ is the bird infectivity rate and *c*_*h*_ is the rate at which a bird gets bitten per unit time. By the conservation of biting property, every bite received by a bird is also a bite taken by a mosquito [27], such that:

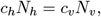

where *N*_*h*_ and *N*_*v*_ represent the total concentration of bird and mosquito populations, respectively. Assuming *N*_*h*_ = *N*_*v*_ = 1, it follows that *c*_*v*_ = *c*_*h*_, and hence:

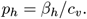

Using the parameter estimates from sequential data fitting, the probability that a bird bitten by an infected mosquito becomes infected with Usutu virus is:

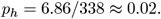

Using the parameter estimates from simultaneous data fitting, the probability that a bird bitten by an infected mosquito becomes infected with Usutu virus is:

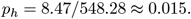

These results suggest that, although simultaneous data fitting yields higher estimates for both the mosquito biting rate and bird infectivity rate, the overall per-bite transmission probability of Usutu virus from mosquito to bird is slightly lower than that obtained from sequential fitting.

### 3.3 Practical Identifiability Results

To address heterogeneity in the data (arising from differences among mosquito lineages or bird species involved in Usutu virus transmission), we assessed the practical identifiability of model parameters by introducing varying degrees of noise into the micro- and macroscale datasets. We fitted parameters sequentially and simultaneously to determine whether uncertainties in the results depend on the chosen data fitting approach.

#### Within-host scale results

We investigated the practical identifiability of the within-host model (Eq. (2.1)) using the Monte Carlo simulation (MCS) method (see Practical Identifiability Analysis section in Materials and Methods for details). A total of 1000 *in-silico* datasets were generated by adding measurement noise (*σ*) to the viral titer data, which were collected every 12 hours over the first 6.5 days (see Table 3). The model was then refitted to each noisy dataset, and MCS was used to compute the average relative estimation error (ARE), thereby quantifying the sensitivity of parameter estimation to measurement noise (see Practical Identifiability Analysis section in Materials and Methods for details). We found that the AREs of all parameters remained below the imposed measurement error for *σ* ∈ {1%, 5%, 10%}, indicating that all model parameters in Eq. (2.1) are strongly practically identifiable under these noise levels (see Definition 2.2 and ARE values in Table 10). Furthermore, for *σ* = 20%, all parameters were found to be weakly practically identifiable (see Definition 2.2 and ARE values in Table 10). These results suggest that, when data is collected every 12 hours, the within-host model remains robust to moderate noise levels, but higher noise diminishes parameter identifiability.

#### Per-bite probability of mosquito infection scale results

Similarly, we evaluated the practical identifiability of the per-bite probability of mosquito infection model (Eq. (2.2)) using the Monte Carlo simulation method. A total of 1000 *in-silico* datasets were generated by adding measurement noise (*σ*) to mosquito infection percent data, which were collected at seven viral titer levels (see Table 4). The model was then refitted to each noisy dataset, and MCS was used to compute the average relative estimation error (ARE), thereby quantifying the sensitivity of parameter estimation to measurement noise (see Practical Identifiability Analysis section in Materials and Methods for details). We found that *h* is weakly practically identifiable and *a* is practically unidentifiable for all *σ* = {1%, 5%, 10%, 20%} noise levels (see Table 11). These results suggest that the per-bite probability of mosquito infection model is not robust to noise, suggesting that the unidentifiable parameter *a* should be removed from data fitting.

#### Between-host scale results

We next evaluated the practical identifiability of the parameters of the between-host model (Eqs. (2.3)–(2.4)), obtained through data fitting to between-host empirical data alone (Table 5), using the Monte Carlo simulation (MCS) method (see Identifiability Analysis in Materials and Methods for details). We found that *β*_*h*_, *c*_*v*_, and *µ*_*v*_ are weakly practically identifiable, whereas *γ*_*h*_ is not practically identifiable at any noise level *σ* = {1%, 5%, 10%, 20%} ( see Definition 2.2 and ARE values in Table 12). These results indicate that the parameters of the between-host model obtained through sequential data fitting are not robust to measurement noise.

#### Multiscale results

Lastly, we evaluated the practical identifiability of the within- and between-host parameters obtained through simultaneous fitting of the multiscale model (Eqs. (2.1)–(2.4)) to virus titer and bird population data. Using the Monte Carlo simulation (MCS) method (see Practical Identifiability Analysis in Materials and Methods for details), we generated 1000 *in-silico* datasets by adding measurement noise (*σ*) to the empirical within- and between-host data (Tables 3 and 5). The multiscale model was refitted to each noisy dataset, and MCS was used to compute the average relative estimation error (ARE), thereby quantifying the sensitivity of parameter estimation to measurement noise. Our results show that all within- and between-host parameters are strongly practically identifiable, with the exception of the bird recovery rate (*γ*_*h*_), which is only weakly identifiable, across all noise levels *σ* = {1%, 5%, 10%, 20%} (see Definition 2.2 and ARE values in Table 13). These findings indicate that parameter estimates obtained through simultaneous fitting across biological scales are robust to measurement noise. Similar improvements in parameter estimation from multiscale data fitting have been reported in previous studies [15, 20].

## 4 Discussion

Using empirical data to quantify unknown parameters in vector-borne dynamical systems is essential for providing meaningful predictions that can guide interventions and robust control [28–33]. By merging mathematical and statistical theory, one can establish appropriate designs for future experiments, improving parameter estimation, consolidating predictions, and revealing model limitations [34]. This problem is inherently complicated when data is collected at different scales of resolution, where feedbacks between scales can propagate uncertainty in quantification [5, 20].

In this study, we addressed the problem of robustness in parameter quantification and model predictions within a multiscale mosquito-borne model of Usutu virus infection in bird and mosquito populations. The multiscale framework integrates two microscale mathematical models, a within-host system describing Usutu virus replication dynamics in birds (modeled using ordinary differential equations) and a probability of mosquito infection (modeled with an algebraic function), into a macroscale between-host transmission model of Usutu virus spread in mosquito and bird populations (formulated using partial differential equations). The multiscale model was validated across these three biological scales using two laboratory-based datasets generated by our group under an optimal experimental design framework [17] and one surveillance-based dataset obtained from the literature [4].

At the individual bird infection scale, we modeled virus–host interactions using within-host models developed for other acute viral infections [35–39], including Usutu virus infection in house sparrows [5] and chickens [10, 11]. We and others have shown that model parameters are uniquely identifiable only when initial conditions are known, even when unlimited data are available [7, 13]. Moreover, our previous work demonstrated that by adding (up to 20%) noise in the data to represent inter-bird variability, the model becomes practically unidentifiable unless data are collected at 12-hour intervals over a 7-day period [5]. To address this, we collected daily data from two cohorts of canary birds, measured 12 hours apart, and combined them to generate a population-level dataset with effective 12-hour sampling intervals [17]. Fitting the within-host model to this dataset, we found that all parameters are identifiable under moderate noise levels (up to 10%), whereas parameter estimates became weakly identifiable when the noise level increased to 20%. These results demonstrate that with appropriately designed sampling strategies and moderate data variability, the within-host viral kinetics model can reliably quantify key parameters of Usutu virus infection dynamics in birds.

At the per-bite probability of mosquito infection scale, we modeled the proportion of mosquitoes infected as a function of the viral titer in their blood meal using a power-law function, as previously described [5]. Prior studies have shown that reliable quantification of the parameters in this function requires identifying the viral concentration threshold that ensures infection in the majority (more than 90%) of exposed mosquitoes. To address this, we conducted a transmission experiment in which mosquitoes fed on cotton balls containing blood meals with viral titers ranging from 3.5 log_10_ to 7.9 log_10_ PFU/ml of the Netherlands 2016 Usutu virus strain [17]. Fitting the power-law model to these data revealed that one parameter (*a*) remained unidentifiable, even under this improved experimental design. Consequently, we fixed this parameter to an arbitrary value and excluded it from further quantitative analysis. These findings indicate that while the power-law framework effectively captures the general relationship between viral titer and mosquito infection probability, full parameter identifiability is not possible under this model formulation.

Finally, at the bird and mosquito population scale, we modeled Usutu virus incidence in wild bird populations using a between-host epidemic model of mosquito–bird interactions, validated against published passive surveillance data on Usutu virus incidence in wild birds collected twice monthly. Because the between-host model incorporates microscale information on viral titers and the per-bite probability of mosquito infection, we employed two strategies for parameter estimation: (i) sequential estimation, in which between-host parameters were estimated from surveillance data alone, after microscale parameters were estimated from microscale data; and (ii) simultaneous estimation, in which both within-host and between-host parameters were estimated jointly using viral titer and surveillance datasets (while unidentifiable parameters from the per-bite probability of mosquito infection model were fixed to fitted values).

Before determining which parameters are quantifiable under noisy conditions, we conducted a structural identifiability analysis of the multiscale model. This analysis revealed that all parameters are uniquely identifiable under ideal (noise-free) conditions, provided that the within-host initial conditions and the initial distribution of infected birds are known. Building on these theoretical results, we next examined practical identifiability under realistic, noisy data scenarios.

Fitting the between-host model to surveillance data alone yielded parameters that were either weakly identifiable or unidentifiable under all noise levels (1–20%). In contrast, simultaneous fitting to both viral titer and surveillance data substantially improved identifiability, with all but one parameter (*γ*_*h*_) becoming strongly identifiable across noise levels of up to 20%. These results demonstrate that simultaneous parameter estimation across biological scales enhances model robustness to measurement noise and improves identifiability. Similar improvements in parameter estimation through multiscale data fitting have been reported in previous studies [15, 20].

In this study, we found that using simultaneous rather than sequential data fitting improved parameter quantification and model predictive performance. We acknowledge, however, that this outcome may be model- and data-dependent. Although simultaneous fitting yielded more robust parameter estimates under noisy conditions, the resulting temporal model predictions, such as viral load dynamics, remained consistent between the two fitting strategies. Importantly, each approach carries distinct advantages and limitations. Sequential fitting simplifies model complexity at each stage but relies on more limited data from individual biological scales. In contrast, simultaneous multiscale fitting integrates a broader and more comprehensive dataset but must contend with data heterogeneity, including differences in magnitude, scale, and sampling frequency across biological levels. When using numerical optimization algorithms (e.g., least squares), this heterogeneity can bias the fitting process toward high-magnitude data, potentially reducing performance for smaller or less frequently sampled datasets [24]. Ultimately, the choice between sequential and simultaneous fitting should be guided by the specific research question, data structure, and modeling objectives.

In conclusion, we developed a multiscale vector-borne model of Usutu virus infection and validated its parameters using both laboratory data collected under an optimally designed experimental framework and published surveillance data from wild bird populations. Using this model, we quantified the robustness of parameter estimates and found that multiscale fitting to integrated datasets improves the reliability and identifiability of model parameters but can lead to slight altering of qualitative model predictions.

## Supporting information

### S1. Numerical Scheme

In this section, we introduce a finite difference method for solving the time-since-infection model Eqs. (2.3)-(2.4). We begin by constructing the numerical mesh by discretizing the domain *D* = {(*t, τ* ) : 0 ≤ *t* ≤ *M*_*t*_, 0 ≤ *τ* ≤ *M*_*τ*_ }, where *τ* is the infection age, *t* is epidemic time, *M*_*t*_ is the maximum time, and *M*_*τ*_ is the maximum infection age. We discretize the spans of infection age and epidemic time into subintervals such that Δ*τ* = *κ*Δ*t*. Thus, the points in the epidemic time and infection age directions are given by *t*_*n*_ = *n*Δ*t* and *τ*_*k*_ = *k*Δ*τ*, respectively, with *M*_*t*_ = *N* Δ*t, M*_*τ*_ = *K*Δ*τ*, and *N* and *K* the total number of subintervals. Let 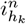 be the approximation of *i*_*h*_ at the point (*t*_*n*_, *τ*_*k*_). We denote the approximate solutions of the state variables at time *t*_*n*_ as 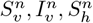, and 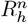. We first discretize the equation for the susceptible mosquitoes, *S*_*v*_, by replacing the time derivative with a backward difference, and obtain:

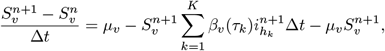

Next, we linearize the nonlinear term by evaluating *i*_*h*_ at epidemic time *t*_*n*_ instead of *t*_*n*+1_, then solve for 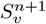 to obtain:

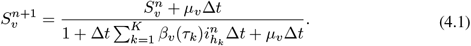

Similarly, we discretize the equations for the infected mosquitoes, *I*_*v*_, and susceptible birds, *S*_*h*_, by evaluating them at *t*_*n*+1_ and applying a backward difference. This process yields the following equations:

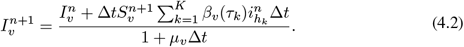

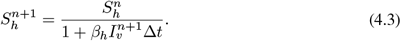

Finally, we discretize the PDE, evaluating it at *t*_*n*+1_ and *τ*_*k*_. We replace the derivative in *τ* with a forward difference and the derivative in *t* with a backward difference. This results in the following difference equation:

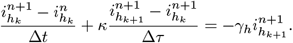

Since 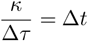,

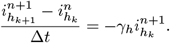

To transform this method into an implicit scheme, we replace the term 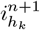 with 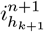, and solve for 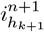 and obtain:

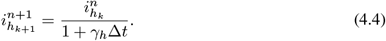

Considering the boundary condition at *t* = *t*_*n*+1_, we have,

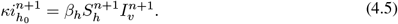

Given that the total bird population *S*_*h*_(*t*) + *I*_*h*_(*t*) + *R*_*h*_(*t*) = 1 remains constant:

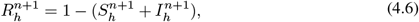

where 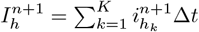. Therefore, the finite difference problem, derived from Eqs. (4.1)-(4.6) becomes:

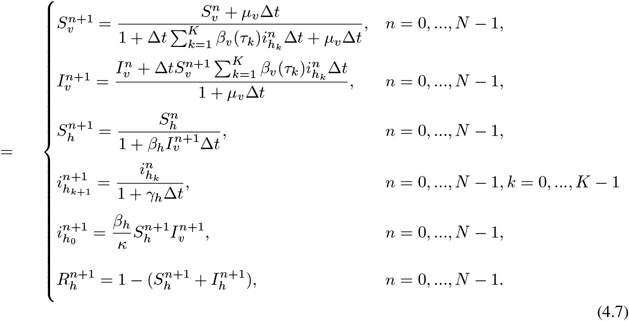

### S2. Monte Carlo simulation results

Practical identifiability results are presented below.

**Table 10.** MCS results for the within-host model: Monte Carlo simulation results for virtual datasets generated at each data point in Table 3. The AREs give the average relative estimation errors for each parameter of the within-host model (Eq. (2.1)) at noise level *σ*.

| Parameter | $\beta$ | $k$ | $\delta$ | $\pi$ | $c$ |
| --- | --- | --- | --- | --- | --- |
| *ARE* |  |  |  |  |  |
| $\sigma = 1\%$ | 0.1 | 0 | 0 | 0 | 0 |
| ----- |  |  |  |  |  |
| *ARE* |  |  |  |  |  |
| $\sigma = 5\%$ | 1.3 | 0.2 | 0.4 | 1.3 | 0.8 |
| ----- |  |  |  |  |  |
| *ARE* |  |  |  |  |  |
| $\sigma = 10\%$ | 3.5 | 0.4 | 1.7 | 3.3 | 1.7 |
| ----- |  |  |  |  |  |
| *ARE* |  |  |  |  |  |
| $\sigma = 20\%$ | 34.6 | 6.5 | 11.1 | 25.9 | 24.9 |

**Table 11.** MCS results for the per-bite percent mosquito infection model: Monte Carlo simulation for virtual datasets generated at each data point in Table 4. The AREs give the average relative estimation errors for each parameter of the probability of infection function Eq. (2.2) at noise level *σ*.

| Parameter | $a$ | $h$ |
| --- | --- | --- |
| $\sigma = 1\%$ | | |
| *ARE* | 10 | 1.6 |
| $\sigma = 5\%$ | | |
| *ARE* | 54.9 | 8.2 |
| $\sigma = 10\%$ | | |
| *ARE* | 119.4 | 17.7 |
| $\sigma = 20\%$ | | |
| *ARE* | 197.1 | 35.2 |

**Table 12.** MCS results for the between-host model: Monte Carlo simulation results for virtual datasets generated at each data point in Table 5. The AREs give the average relative estimation errors for each parameter of the between-host model given in Eqs. (2.3)-(2.4) at noise level *σ*.

| Parameter | $\beta_h$ | $c_v$ | $\gamma_h$ | $\mu_v$ |
| --- | --- | --- | --- | --- |
| *ARE* |  |  |  |  |
| $\sigma = 1\%$ | 2.8 | 4.1 | 16.8 | 1.2 |
| *ARE* |  |  |  |  |
| $\sigma = 5\%$ | 16.9 | 31 | 75.7 | 6.3 |
| *ARE* |  |  |  |  |
| $\sigma = 10\%$ | 30.4 | 65.1 | 136.5 | 10.7 |
| *ARE* |  |  |  |  |
| $\sigma = 20\%$ | 51.1 | 99.1 | 166.6 | 18.5 |

**Table 13.** MCS results for the multiscale model: Monte Carlo simulation results for virtual datasets generated at each data point in Tables 3 and 5. The AREs give the average relative estimation errors for each parameter of the multiscale model given in Eqs. (2.1)-(2.4) at noise level *σ*.

| Parameter | $\beta_h$ | $c_v$ | $\gamma_h$ | $\mu_v$ | $\beta$ | $k$ | $\delta$ | $\pi$ | $c$ |
| --- | --- | --- | --- | --- | --- | --- | --- | --- | --- |
| *ARE* |  |  |  |  |  |  |  |  |  |
| $\sigma = 1\%$ | 0.4 | 0 | 16.8 | 0.6 | 0.9 | 2.3 | 3.9 | 13.5 | 4.2 |
| *ARE* |  |  |  |  |  |  |  |  |  |
| $\sigma = 5\%$ | 1.5 | 0 | 47.8 | 2.4 | 3.2 | 0.6 | 1.4 | 6 | 2 |
| *ARE* |  |  |  |  |  |  |  |  |  |
| $\sigma = 10\%$ | 4.4 | 0 | 84.2 | 6.3 | 7.5 | 0.7 | 1.7 | 9.9 | 3.6 |
| *ARE* |  |  |  |  |  |  |  |  |  |
| $\sigma = 20\%$ | 12.6 | 0 | 146.3 | 14.1 | 17.7 | 2.7 | 4 | 19 | 7.1 |

## Acknowledgments

Funding for this work was provided by NIH NIGMS 1R01GM152743. SMC and QM acknowledge partial support from National Science Foundation (NSF) grant No. 2051820. This research was enabled in part through the Virginia Tech Center for the Mathematics of Biosystems (VT-CMB).

## Author Contribution

Conceptualization: NT, ND, SMC; Formal analysis: NT, YRL, QMM, SMC; Data analysis: RP, ND; Funding acquisition: NT, ND, SMC; Software: NT, YRL, QMM; Supervision: NT, ND, SMC; Writing – original draft: NT, SMC; Writing – review & editing: NT, YRL, QMM, RP, ND, SMC.

